# Differential representations of spatial location by aperiodic and alpha oscillatory activity in working memory

**DOI:** 10.1101/2025.03.21.644412

**Authors:** Andrew Bender, Chong Zhao, Edward Vogel, Edward Awh, Bradley Voytek

## Abstract

Decades of research have shown working memory (WM) relies on sustained pre-frontal cortical activity and visual extrastriate activity, particularly in the alpha (8-12 Hz) frequency range. This alpha activity tracks the spatial location of WM items, even when spatial position is task-irrelevant and there is no stimulus currently being presented. Traditional analyses of putative oscillations using bandpass filters, however, conflate oscillations with non-oscillatory aperiodic activity. Here, we reanalyzed seven different human electroencephalography (EEG) visual WM datasets to test the hypothesis that aperiodic activity–which is thought to reflect the relative contributions of excitatory and inhibitory drive–plays a distinct role in visual WM from true alpha oscillations. To do this, we developed a novel, time-resolved spectral parameterization approach to disentangle oscillations from aperiodic activity during WM encoding and maintenance. Across all seven tasks, totaling 112 participants, we captured the representation of spatial location from total alpha power using an inverted encoding model (IEM), replicating traditional analyses. We then trained separate IEMs to estimate the strength of spatial location representation from aperiodic-adjusted alpha (reflecting just the oscillatory component) and aperiodic activity, and find that IEM performance improves for aperiodic-adjusted alpha compared to total alpha power that blends the two signals. We also discover a novel role for aperiodic activity, where IEM performance trained on aperiodic activity is highest during stimulus presentation, but not during the WM maintenance period. Our results emphasize the importance of controlling for aperiodic activity when studying neural oscillations while uncovering a novel functional role for aperiodic activity in the encoding of visual WM information.

**Significance statement:** Working memory is a crucial component of cognition, yet its neural mechanisms are not fully understood. Research shows that alpha activity – presumed to reflect neural oscillations – tracks the location of items we hold in memory. However, these analyses assume that all alpha power is oscillatory, even though oscillations are mixed with non-oscillatory, aperiodic activity that may be physiologically and functionally distinct. Here, we use a novel analytical approach for separating alpha oscillations and aperiodic activity dynamically across time. Our results reveal distinct roles for each in human visual working memory: aperiodic activity encodes the spatial location of information whereas alpha oscillations maintain the location of that information.

## Introduction

Working memory (WM) refers to our ability to hold and manipulate information in mind for a short time, usually just a few seconds. Although WM is core to many cognitive functions, and is disrupted in numerous psychiatric and neurological disorders, the underlying neurocomputational principles of WM remain elusive. While sustained single-neuron activity in the prefrontal cortex (PFC) is considered a hallmark of WM, it relies on population coding distributed across many regions (Christophel, Klink, et al., 2017). Population neural activity in the PFC represents a broad array of task variables such as stimulus-stimulus association (Stokes et al., 2013), and sustained representations of WM content in the pattern of activity of voxels distributed within parietal and sensory cortices (Christophel, Hebart, and Haynes, 2012; Harrison and Tong, 2009; Riggall and Postle, 2012; Serences et al., 2009; Sreenivasan, Vytlacil, and D’Esposito, 2014).

Another way of assessing population codes is via extracellular field recordings that measure synaptic and transmembrane currents in a neural population (Buzsáki, Anastassiou, and Koch, 2012). Oscillations are a common feature of these signals, and play key roles in neural communication and computation (Fries, 2005). Changes in oscillatory power have been documented in the delay period of WM tasks at multiple scales: from invasive local field potentials (LFPs) to noninvasive electroencephalography (EEG) and magnetoencephalography (MEG). Visual cortical alpha (8–12 Hz) oscillations have been implicated in WM, with both alpha power increases (Jensen, 2002; Jokisch and Jensen, 2007; Klimesch, 2012) and decreases (Jokisch and Jensen, 2007; Ede, Jensen, and Maris, 2017; Adam, Robison, and Vogel, 2018; Van Engen et al., 2024) during WM tasks. These seemingly paradoxical results have been resolved by differential alpha dynamics, with increased alpha power reflecting disengagement of task-irrelevant sensory representations, and decreased alpha power reflecting engagement of task-relevant sensory representations (Jokisch and Jensen, 2007; Spitzer and Blankenburg, 2012; Ede, 2018). Like its role in spatial attention, alpha power is reduced contralateral to locations held in spatial WM (Medendorp et al., 2007; Van Der Werf et al., 2008; Dijk et al., 2010), with both healthy aging and unilateral PFC lesions altering visual cortical delay period alpha (Tran, Hoffner, et al., 2016; Voytek and Knight, 2010). Alpha band activity also encodes the precise spatial location of WM stimuli (Bae and Luck, 2018; Foster, Sutterer, et al., 2016), even when spatial location is task-irrelevant (Foster, Bsales, et al., 2017).

While there is substantial evidence for changes in narrowband alpha power in visual attention and WM, oscillations occur in conjunction with non-oscillatory aperiodic activity. Aperiodic activity has historically been dismissed as neural noise, though given the physiological nature of the LFP (and thus, by extension, EEG), more recent work has shown that aperiodic activity likely reflects asynchronous cortical drive that is coarsely related to the relative excitatory/inhibitory drive (Chini, Pfeffer, and Hanganu-Opatz, 2022; Gao, Peterson, and Voytek, 2017), and is dynamically related to cognitive and behavioral state Tran, Rolle, et al., 2020; Voytek, Kramer, et al., 2015; Waschke, Wöstmann, and Obleser, 2017. Critically, changes in aperiodic activity are distinct from event-related potentials (Arnett, Peisch, and Levin, 2022), suggesting that event-related fluctuations in aperiodic activity are not merely a byproduct of phase-locked responses but instead reflect broadband shifts in neural excitability. Using traditional spectral analyses, such as narrowband filtering in a predefined alpha band, dynamic and task-related changes in this aperiodic activity can appear to be changes in oscillatory activity, even if no oscillation is present (Donoghue, Haller, et al., 2020; Donoghue, Schaworonkow, and Voytek, 2021). This raises the question as to whether the preponderance of evidence of alpha dynamics during WM reflects true alpha oscillations, or is actually capturing rapid fluctuations in aperiodic dynamics that manifest as power changes in the alpha frequency range, or reflects dynamic changes in both.

Here, we reanalyze seven previously published EEG experiments, totalling 112 participants, that have previously demonstrated alpha band encoding of visual WM information during a WM delay period under various task conditions. By using a novel, time-resolved extension of spectral parameterization, we dissociate the effects of event-related alpha oscillations from aperiodic dynamics in the encoding and maintenance of WM information with near-millisecond resolution. Even when controlling for aperiodic dynamics, we find that alpha power represents the correct spatial location of visual items sustained across the WM delay period across all seven tasks. Importantly, this representation actually improves after controlling for aperiodic activity. This reinforces the idea that alpha oscillations – not just alpha power – are critical for maintaining information in visual WM. We also discover a novel role of aperiodic activity in the transient encoding of spatial information across all seven tasks, that, unlike alpha oscillations, does not sustain during the delay period but is only evident after stimulus presentation. These results highlight the importance of considering aperiodic activity in our analyses. They strengthen the evidence for the role of alpha oscillations in visual WM while hinting at a previously unknown role of aperiodic activity, potentially for the encoding of visual information.

## Materials and Methods

### Participants

The EEG data analyzed in this paper was collected from participants at the University of Oregon (*n* = 56) and the University of Chicago (*n* = 56). These data are from three publicly available datasets (Foster, Sutterer, et al., 2016; Foster, Bsales, et al., 2017; Sutterer et al., 2019), and the published reports contain the complete details on participants, data collection procedures, experimental design, and data preprocessing. But we summarize them briefly, here.

### Experimental design

The three publicly available datasets (Foster, Sutterer, et al., 2016; Foster, Bsales, et al., 2017; Sutterer et al., 2019) encompass data from seven different tasks/experiments (Table 1), during all of which participants attended to and remembered positions of memory items around fixation, across a delay period. As shown in Fig. 1, the tasks proceeded in three stages: stimulus, delay, and response. During stimulus presentation, participants were presented with one or two simple stimuli (circles, squares, gratings, or triangles) equidistant from fixation (4° visual angle). Stimuli were presented for durations ranging from 100 ms to 1000 ms. Following stimulus offset, there was a fixed, blank delay period containing only a fixation point where durations ranged from 1000–1750 ms depending on the dataset. Finally, during the response period, participants were presented with a response screen that contained either a ring presented around fixation (continuous recall tasks) or a single probe stimulus (change detection task). All of the tasks were continuous recall tasks except for Task 3, which was a change detection task in which the participants had 250 ms to determine whether the probe stimulus had changed from stimulus presentation.

**Table 1:**
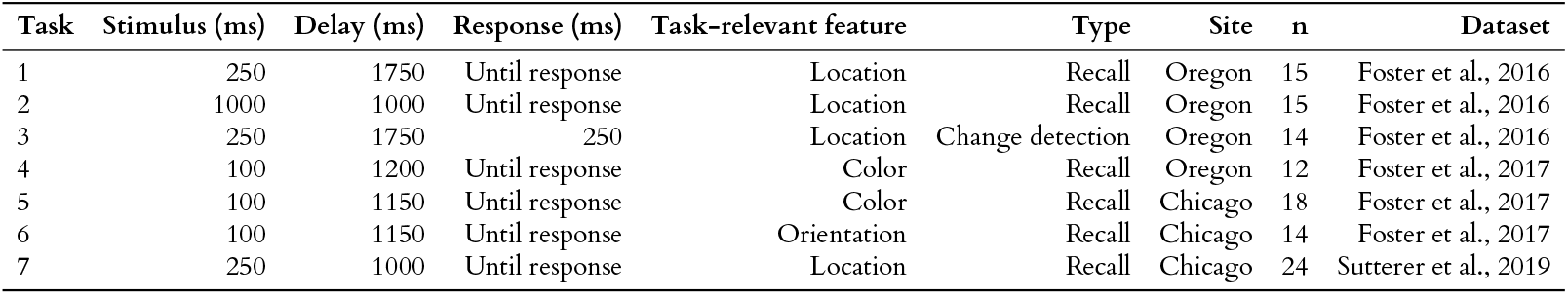
Task parameters across seven WM tasks and two research sites. Stimulus presentation and delay timings, response type, recording site, and which feature was relevant for the WM task were variable across the tasks.

| Task | Stimulus (ms) | Delay (ms) | Response (ms) | Task-relevant feature | Type | Site | n | Dataset |
| --- | --- | --- | --- | --- | --- | --- | --- | --- |
| 1 | 250 | 1750 | Until response | Location | Recall | Oregon | 15 | Foster et al., 2016 |
| 2 | 1000 | 1000 | Until response | Location | Recall | Oregon | 15 | Foster et al., 2016 |
| 3 | 250 | 1750 | 250 | Location | Change detection | Oregon | 14 | Foster et al., 2016 |
| 4 | 100 | 1200 | Until response | Color | Recall | Oregon | 12 | Foster et al., 2017 |
| 5 | 100 | 1150 | Until response | Color | Recall | Chicago | 18 | Foster et al., 2017 |
| 6 | 100 | 1150 | Until response | Orientation | Recall | Chicago | 14 | Foster et al., 2017 |
| 7 | 250 | 1000 | Until response | Location | Recall | Chicago | 24 | Sutterer et al., 2019 |

**Figure 1:**
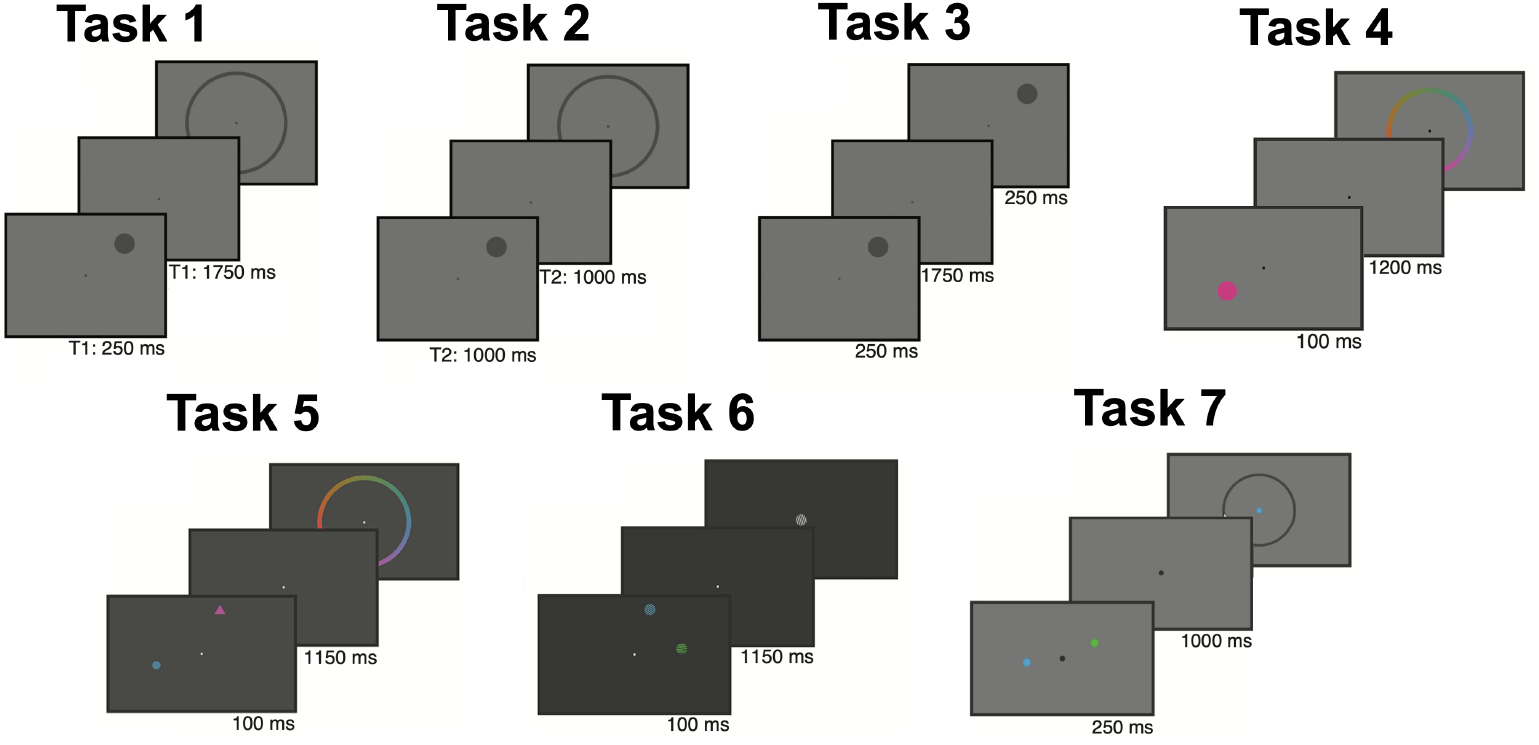
Example task for each WM task. In each task, a stimulus was presented (front), the participant was asked to remember a feature of the stimulus across a WM delay (middle), and then reported the remembered feature value during a response period (back).

Importantly, all tasks required that participants attend to locations around fixation either explicitly or implicitly, regardless of the response required. In the University of Oregon sample, participants were instructed to remember the spatial position (Tasks 1–3, *n* = 44) or color (Task 4, *n* = 12) of a circle stimulus presented around fixation throughout the blank delay period. In the University of Chicago sample, participants were presented with two stimuli (a square and a triangle in Task 5, two colored gratings in Task 6, and two colored circles in Task 7) and cued in advance to attend and remember only one of the stimuli based on its shape (Task 5) or color (Tasks 6 and 7). In Task 7, some of the trials featured only one colored circle and no distractor. After a brief delay, participants were required to report the orientation (Task 5), color (Task 6), or spatial position (Task 7) of the cued stimulus. Notably, all tasks required that participants attend spatial positions around fixation and encouraged participants to maintain those spatial representations through a blank delay.

### Spectral decomposition and time-resolved parameterization

Repeating the procedure from the original papers (Foster, Sutterer, et al., 2016; Foster, Bsales, et al., 2017; Sutterer et al., 2019), alpha total power was computed for each trial using a narrowband alpha filter and Hilbert transform approach. The data were filtered in the alpha band (8–12 Hz), and then the Hilbert transform was applied on the filtered data to get the analytic signal. The instantaneous alpha power was the magnitude of this analytic signal. This measure of instantaneous alpha power does not account for confounding contributions of aperiodic activity, and, as such, is hereafter referred to as total alpha power to disambiguate it from our subsequent analyses that separate putative oscillations from aperiodic activity.

To estimate an aperiodic-adjusted instantaneous measure of alpha power, we first computed sliding-window power spectra for each trial using multitapers in MNE-Python (Gramfort, 2013) with time windows of 1 second (Fig. 2B). This procedure resulted in power estimates for 128 frequencies, from 2-50 Hz, from which the “instantaneous” spectral parameters could be estimated. It is important to emphasize that “instantaneous” analytics amplitude using the traditional filter-Hilbert approach has similar temporal uncertainty as our sliding-window short-time Fourier-multitaper approach (Bruns, 2004). Spectral decomposition of these time-frequency representations was then done using the spectral parameterization method (version 1.1.0) (Fig. 2C) (Donoghue, Haller, et al., 2020). Settings for the algorithm were set as: peak width limits: (2, 8); max number of peaks: 4; minimum peak height: 0; peak threshold: 2.0; and aperiodic mode: ‘fixed’. Power spectra were parameterized across the frequency range 2 to 50 Hz. We extracted the aperiodic exponent from these model fits, providing an estimate of the “instantaneous” aperiodic exponent for each time point in each trial (Fig. 2D, orange line). Finally, we estimated the instantaneous aperiodic-adjusted alpha power by subtracting the aperiodic activity from the power spectra for each time point and calculated the area under the curve (AUC) on a linear scale in the alpha range (8–12 Hz) (Fig. 2C). This provided a continuous estimate of alpha oscillatory power (not total alpha power) for each time point in each trial (Fig. 2D, dark purple line). These sliding-window estimates of the aperiodic exponent and alpha oscillatory power were then used with the alpha total power in subsequent analyses.

**Figure 2:**
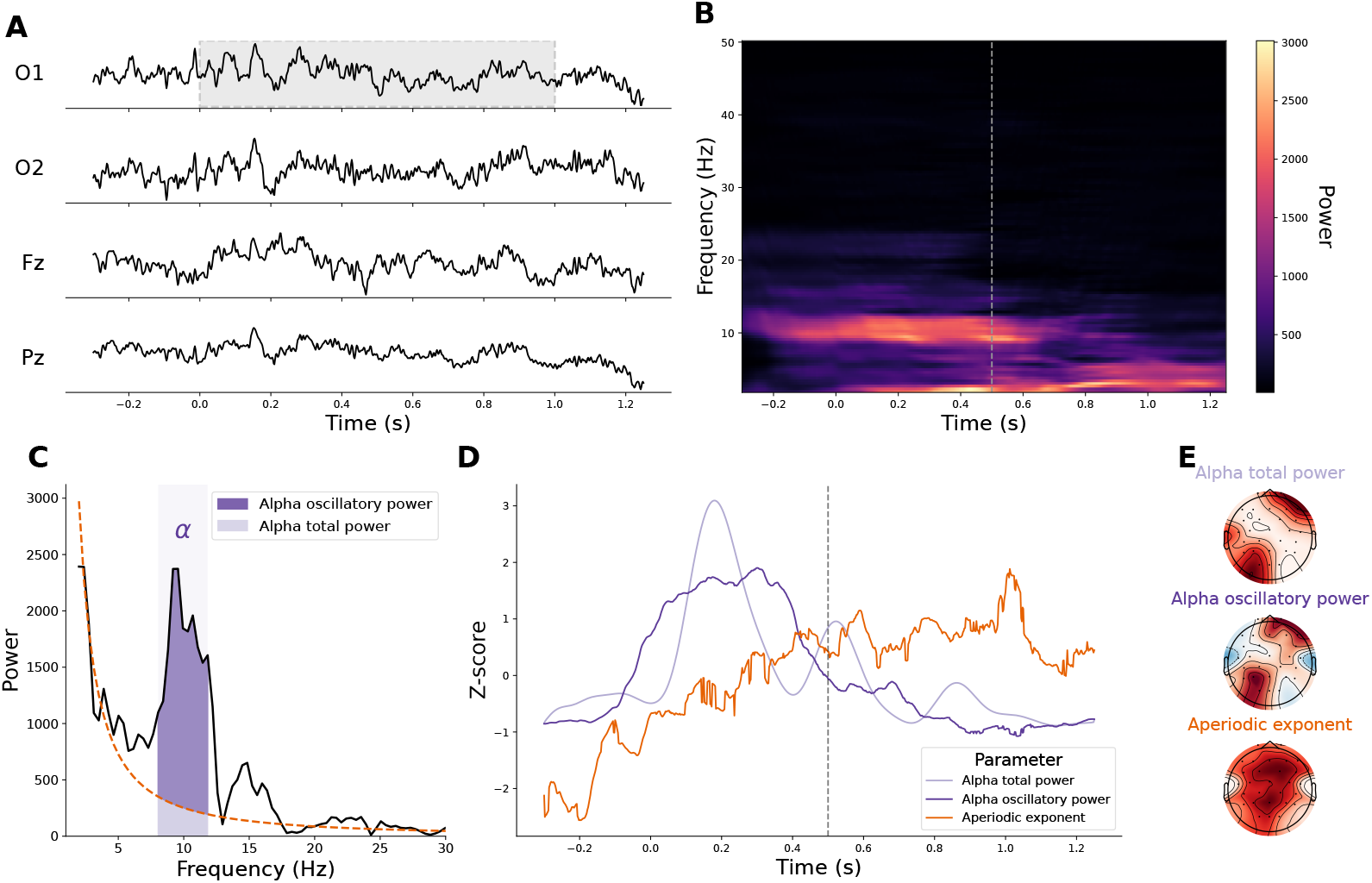
Estimation of time-resolved aperiodic and aperiodic-adjusted oscillatory parameters for individual trials. (A) Voltage traces from one trial for two occipital channels (O1, O2), one parietal channel (Pz), and one frontal channel (Fz). The gray bounding box highlights the one-second temporal window that serves as input for multitaper decomposition to estimate the spectral properties for the time point 0.5 seconds after stimulus presentation. (B) Multitaper spectrogram for one occipital channel (O1) for the same trial as shown in (A). The vertical grey dashed line denotes the time point 0.5 seconds after stimulus presentation. (C) Power spectrum for the time window and channel denoted in (B), which is used to estimate the “instantaneous” spectral parameters. Alpha total power (hashed window) is the full area under the curve (AUC) within the alpha band (8–12 Hz, light purple), while alpha oscillatory power is the alpha total power minus the power attributed to the aperiodic exponent (dark purple). (D) Time-resolved estimates of alpha total power, alpha oscillatory power, and the aperiodic exponent for the same trial as shown in (A) for channel O1. (E) Spatial distribution of alpha total power, alpha oscillatory power, and aperiodic exponent for same time point and trial shown in (A).

To increase the signal-to-noise of these sliding-window estimates, the data were averaged across trials into three separate blocks. Trials were randomly assigned to one of the three blocks, but the same number of trials from each location bin were included in each block and the blocks were independent (i.e., no trial was included in multiple blocks). The values were then averaged across each block, such that the final matrix of parameter values was *l* location bins * *b* blocks × *m* channels × *s* time points.

### Fitting inverted encoding models

Similar to the procedure from the original papers (Foster, Sutterer, et al., 2016; Foster, Bsales, et al., 2017; Sutterer et al., 2019), we fit inverted encoding models (IEMs) to reconstruct location-selective channel tuning functions (CTFs) from the topographic distribution of each parameter (alpha and aperiodic) across electrodes. This was done separately for alpha total power values from the Hilbert transform and the aperiodic exponent and alpha oscillatory power values from spectral parameterization. We assumed that the parameter value at each electrode reflects the weighted sum of eight spatially-tuned channels (i.e., neuronal populations), each tuned for a different angular location (Foster, Sutterer, et al., 2016; Sprague, Ester, and Serences, 2014). Each spatial channel’s response across angular locations was modeled as a half sinusoid raised to the seventh power, given by:

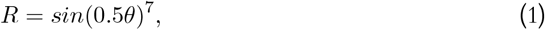

where *θ* is angular location (ranging from 0° to 359°), and *R* is the response of the spatial channel in arbitrary units. This response profile was circularly shifted for each channel such that the peak response of each spatial channel was centered over one of the eight location bins (i.e., 0°, 45°, 90°, etc.). The predicted channel responses for each location bin were derived from these basis functions (calculated using the angular location at the center of each bin).

First, encoding models were fit separately to the alpha total power, aperiodic exponent, and alpha oscillatory power values for each time point (Fig. 3A-D). Data from the first two blocks were used for training, while the third was held-out for testing (described below). Let **B**_**train**_ (*m* electrodes × *n*_*train*_ observations) be the parameter value at each electrode for each measurement in the training set (Fig. 3C), with **C**_**train**_ (*k* channels × *n*_*train*_ observations) being the predicted response of each spatial channel (determined by the basis functions) for each measurement (Fig. 3B), and **W** (*m* electrodes × *k* channels) being a weight matrix that characterizes a linear mapping from “channel space” to “electrode space”. The relationship between **B**_**train**_, **C**_**train**_, and **W** can be described by a general linear model of the form:

**Figure 3:**
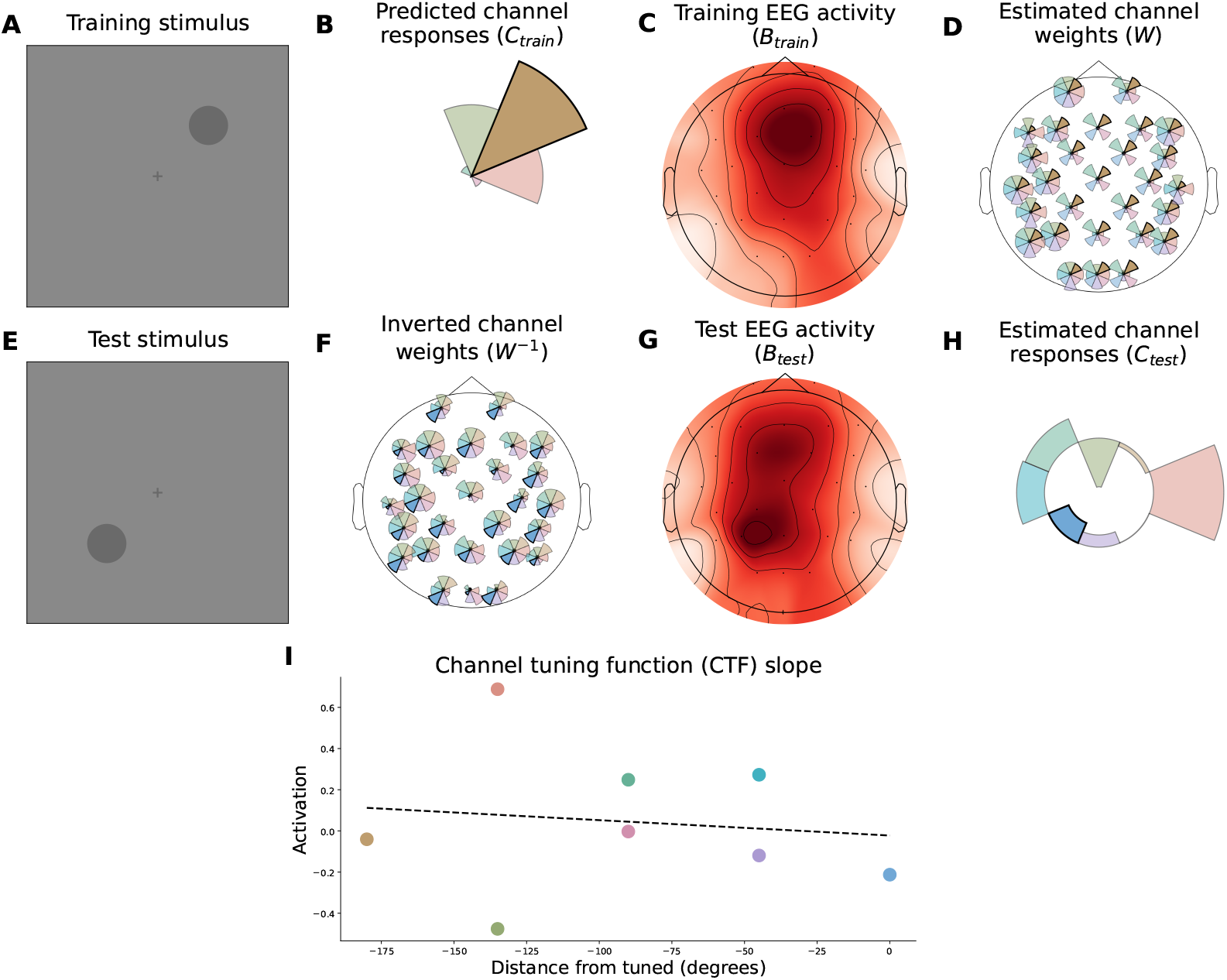
Procedure for fitting inverted encoding models to estimate strength of representation for correct spatial location. (A) Training stimulus presented at 45°. (B) Predicted spatial channel response for neural populations encoding each of the eight location bins for the stimulus presented at 45°. The spatial channel response is modeled using Eq. (1). This corresponds to one row of **C**_**train**_. (C) Average EEG activity for each sensor for one example training block. The topomap shown is the average, aperiodic-adjusted alpha oscillatory power at 0.5 seconds for each sensor for one participant. Each row in **B**_**train**_ represents alpha oscillatory power at 0.5 seconds across each sensor for each trial. (D) The estimated channel weights fit from the predicted channel responses and alpha oscillatory power using Eq. (3). The weights corresponding to the training stimulus shown at 45° are highlighted for each sensor. (E) Test stimulus presented at 225°. (F) Inverted channel weights as shown in Eq. (4). The weights corresponding to the test stimulus shown at 225° are highlighted for each sensor. (G) Average EEG activity for each sensor for one example test block. The topomap shown is the average alpha oscillatory power at 0.5 seconds for each sensor for one participant. Each row in **B**_**test**_ represents alpha oscillatory power at 0.5 seconds across each sensor for each trial. (H) Estimated spatial channel response for neural populations encoding each of the eight location bins for a stimulus presented at 225°, as calculated by Eq. (4). (I) Channel tuning function (CTF) slope, calculated after the spatial estimated channel response shown in (H) is circularly shifted to the correct spatial location and reflected about that spatial location. The dotted line shows the slope of the linear regression of this CTF for stimuli presented at 225° for this test block.

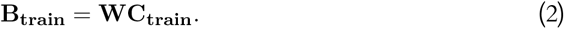

The weight matrix was fit to the training data using least-squares estimation:

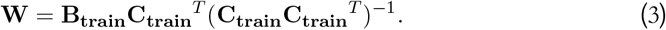

Second, the weights were then inverted and used to transform the test data **B**_**test**_ (*m* electrodes × *n*_*test*_ observations) into estimated channel response **C**_**test**_ (*k* channels × *n*_*train*_ observations, Fig. 3F):

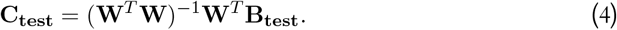

Each estimated channel response function was circularly shifted to a common center by aligning the estimated channel responses to the channel tuned for the stimulus bin to yield CTFs (Fig. 3H). This procedure of fitting an encoding model on the training data **B**_**train**_ and inverting that encoding model to estimate the channel responses for the test data **B**_**test**_ was repeated for each time point and with each block serving as the test block, such that we performed a “leave-one-out” cross-validation. The resulting CTFs were averaged across the three testing blocks. Finally, the procedure was repeated for 100 block assignments for each participant to minimize the influence of idiosyncrasies in estimates of parameter values specific to certain assignments of trials to blocks.

### Statistical analyses

CTFs were circularly shifted to the correct spatial location and reflected about that spatial location, such that points equidistant from the correct spatial location are treated identically.

To estimate the representation strength of the correct spatial location, the slope of the linear regression for these transformed CTFs was calculated, hereafter referred to as the CTF slope (Fig. 3I). We *z*-scored the CTF slopes for each participant based on the baseline period, such that CTF slope values above 1.96 indicate *p* < 0.05. We computed one-sample *t*-tests to evaluate whether CTF slopes in the encoding and delay periods differed significantly from the baseline (which would have a *z*-scored CTF slope of 0 by design). Comparisons between parameters (e.g., alpha total power and alpha oscillatory power) were made using paired *t*-tests across participants. We accounted for multiple comparisons using the Benjamini–Hochberg procedure (Benjamini and Hochberg, 1995).

## Results

### Time courses of spatial location representation are consistent across seven different working memory tasks

We reconstructed the time courses of spatial location representation by alpha total power, alpha oscillatory power, and the aperiodic exponent for each of the seven WM tasks in the composite dataset (Fig. 4). Importantly, these tasks had different stimulus durations, task-relevant features, and were collected across two different sites. Yet, the time courses of spatial location representation by these features were consistent across tasks, with the spatial location representation by alpha total power and alpha oscillatory power peaking during the delay period and the spatial location representation by aperiodic activity peaking during the encoding period.

**Figure 4:**
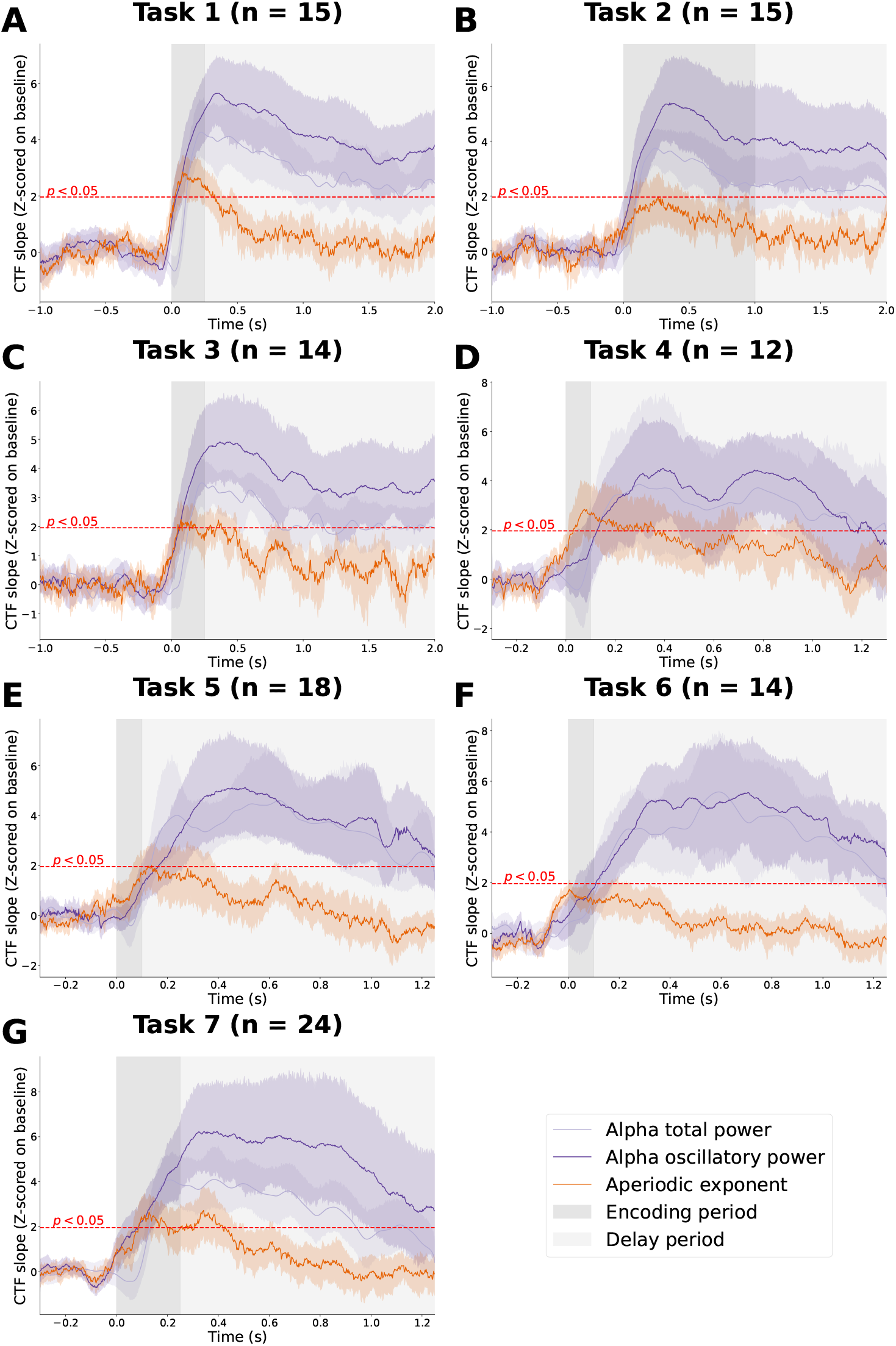
Representation strength of correct spatial location across seven distinct working memory tasks. (A-G) Channel tuning function (CTF) slope, *z*-scored on the baseline period, across each of the seven WM tasks for spatial location by alpha total power (light purple), alpha oscillatory power (dark purple), and the aperiod1i0c exponent (orange). The shading of lines reflect the 95% confidence intervals across participants within each task. The red dotted line indicates *z*-scores that correspond to *p* < 0.05. The darker shading designates the encoding period, while the lighter shading designates the delay period.

### Alpha power represents the correct spatial location during the delay period

We performed one-sample *t*-tests on the *z*-scored CTF slopes across participants for each task. The representation of the correct spatial location during the delay period by alpha total power and by alpha oscillatory was significant across all seven tasks (Fig. 5A and Fig. 5B, respectively). The representation of the correct spatial location during the encoding period by alpha total power was also significant for four of the seven tasks, while that by alpha oscillatory was significant for five of the seven tasks.

**Figure 5:**
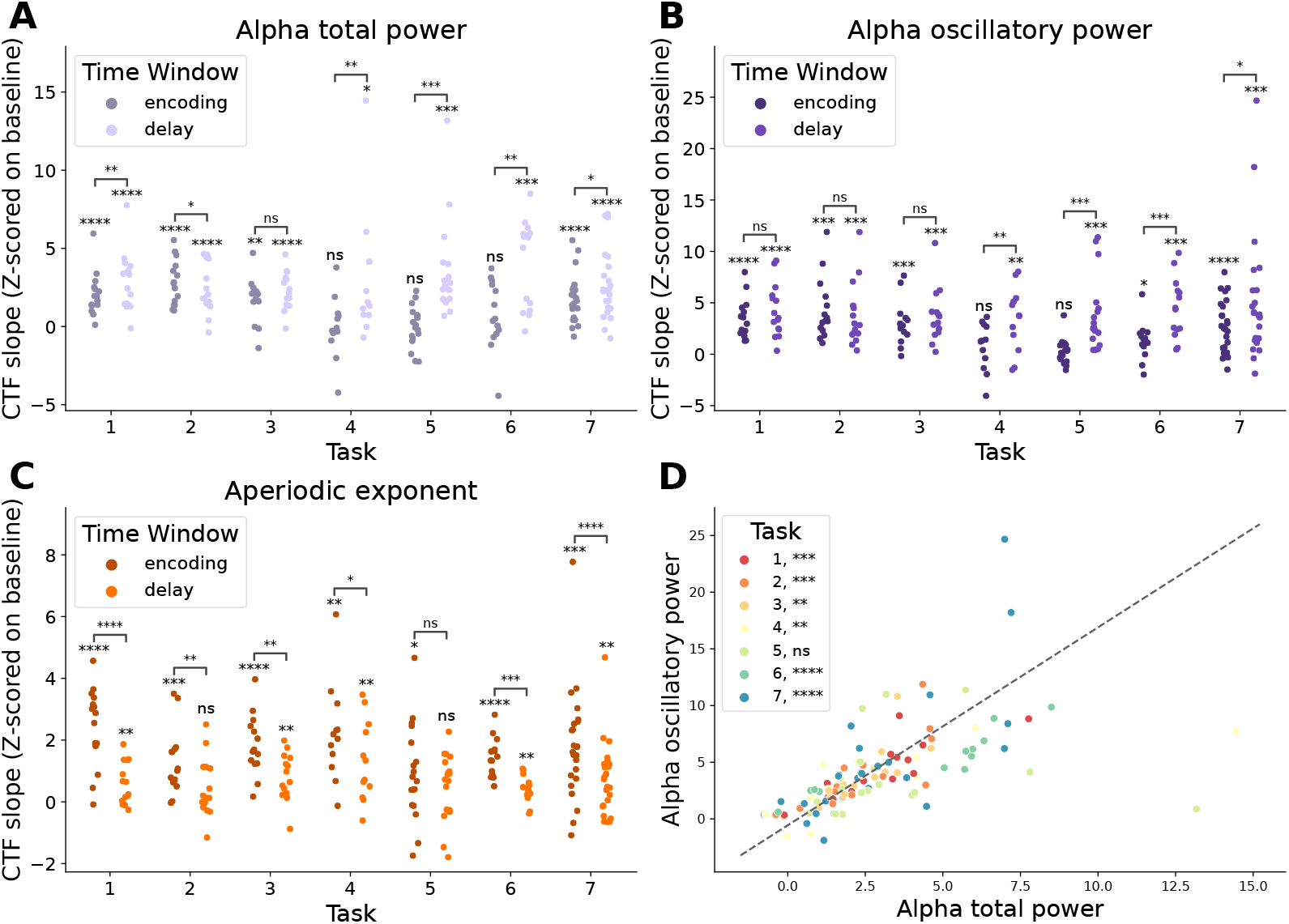
Comparison of representation strength time courses by periodic and aperiodic parameters. (A) Representation strength during encoding and delay periods for each participant by alpha total power, (B) alpha oscillatory power, and (C) aperiodic exponent. Significance above each cluster reflects the deviation from baseline for each time period, while the significance between clusters reflects the paired difference between the encoding and delay time periods across participants. (D) Comparison of representation strength by alpha oscillatory power and alpha total power during the delay period for each participant. Values above the diagonal indicate stronger representation of the correct spatial location by alpha oscillatory power than alpha total power. The significance of the paired *t*-test for each participant’s representation strength of the correct spatial location by alpha oscillatory power versus that by alpha total power across each task is shown in the legend.

To determine whether the strength of representation was significantly different between the encoding and delay periods, we also performed paired *t*-tests between the *z*-scored CTF slopes during the encoding and delay periods. Alpha total power represented the correct spatial location significantly more strongly during the delay, compared to the encoding period, in four of seven tasks, while the encoding period representation was stronger compared to the delay in one task (Fig. 5A). Alpha oscillatory power represented the correct spatial location significantly more strongly during the delay, compared to the encoding period, in four tasks; in the other three tasks, the difference between the encoding and delay period was not significant (Fig. 5B).

### Aperiodic exponent represents the correct spatial location during the encoding period

The representation of the correct spatial location during the encoding period by aperiodic exponent was significant across all seven tasks (Fig. 5C). The aperiodic exponent showed significant representation of the correct spatial location during the delay period in five of the seven tasks. Finally, the representation of the correct spatial location by the aperiodic exponent was significantly stronger during the encoding, compared to the delay period, in six of the seven tasks.

### Alpha oscillatory power better represents the correct spatial location during the delay period than alpha total power

Alpha oscillatory power represented the correct spatial location better than alpha total power during the delay in 85 out of 112 participants (Fig. 5D). To test whether the representation of the correct spatial location by alpha oscillatory power was significantly stronger than that by alpha total power, we computed paired *t*-tests between each participant’s alpha oscillatory power and alpha total power CTF slopes within each task, correcting for multiple comparisons across tasks. The representation of the correct spatial location was significantly stronger in six of the seven tasks (Fig. 5D, legend), suggesting that alpha oscillatory power better captures spatial location information than alpha total power.

## Discussion

In this paper, we applied novel, time-resolved spectral parameterization to estimate continuous aperiodic-adjusted alpha power (i.e., alpha oscillatory power) and aperiodic activity across a composite dataset comprising 112 participants and seven different WM tasks. We first replicated previous findings of significant representation of the correct spatial location by alpha total power during the delay period. After adjusting for contributions of aperiodic activity to these alpha power estimates, we showed significant improvements in the strength of spatial location representation. Finally, we uncovered a novel role for aperiodic activity within the context of working memory–the aperiodic exponent encodes spatial location during stimulus presentation. Together, these results suggest alpha oscillations and aperiodic activity play distinct roles in the encoding and retention of spatial location during working memory.

### Implications

The fact that alpha oscillatory power represents the correct spatial location significantly more strongly than alpha total power in six of the seven tasks shows that traditional analysis approaches that conflate oscillations with aperiodic activity are impairing our ability to understand the functional role of neural oscillations while also hiding the potentially distinct functional role of aperiodic activity. Our results show that alpha oscillatory power better captures the processes that underlie the retention of spatial location in visual WM. These results highlight the importance of removing aperiodic contributions to measures of oscillatory activity, even if aperiodic activity is not the focus of investigation. Apart from its role in WM, differences in alpha total power have been observed in the context of many cognitive functions and cognitive disorders (Başar et al., 1997; Klimesch, Sauseng, and Hanslmayr, 2007; S Palva and JM Palva, 2007; Uhlhaas et al., 2008). In these studies, researchers most commonly use measures of alpha power that do not account for aperiodic activity. Our results across these seven WM tasks demonstrate that using an aperiodic-adjusted measure of alpha power may provide a more sensitive measure of alpha oscillatory activity. Moreover, there may be interesting differences in aperiodic activity that coincide with those already observed in alpha total power.

Aperiodic activity has been shown to capture information about its underlying physiological generators, such as the relative balance of excitation and inhibition (Gao, Peterson, and Voytek, 2017). Moreover, small, dynamic fluctuations in excitation-inhibition balance are critical for healthy cognitive functioning, allowing for efficient information transmission and gating (TP Vogels and LF Abbott, 2009), network computation (Mariño et al., 2005), top-down attentional gain modulation (Chance, L Abbott, and Reyes, 2002; Kaliukhovich and R Vogels, 2016; Reynolds and Heeger, 2009), and WM maintenance(Lim and Goldman, 2013). The model of aperiodic activity developed by Gao and colleagues (Gao, Peterson, and Voytek, 2017) suggests that an increase in excitation necessary for encoding the stimulus would manifest as a decrease in the aperiodic exponent, leading to a flattening of the power spectrum. Recent work has shown a flattening of the power spectrum in intracranial EEG during memory encoding that is consistent with this hypothesis (Preston, Schaworonkow, and Voytek, 2025). Our result that the aperiodic exponent represents the correct spatial location during the encoding period is consistent with these previous findings and further validates the notion that the dynamic changes in the aperiodic exponent reflect increases in excitation necessary for stimulus encoding. Future studies are necessary to test directly whether an increase in excitation underlies the representation of spatial location by aperiodic exponent during encoding. Such studies would validate both the computational model of aperiodic activity as reflective of relative balance in excitation and inhibition and the role of excitation-inhibition interactions in encoding stimulus for WM.

Our results demonstrate that the aperiodic exponent represents spatial location during the encoding period, while alpha oscillatory activity represents spatial location during the delay period. Previous work has demonstrated that E-I interactions are essential for the formation of neural oscillations (Atallah and Scanziani, 2009; Sohal et al., 2009), which suggests that oscillations and aperiodic activity interact.

### Limitations

While alpha oscillatory activity more strongly represents spatial location than alpha total power in all but one of the seven WM tasks tested here, it is more computationally costly to do sliding-window spectral parameterization than to use traditional methods like a filter-Hilbert approach. That said, there are a couple of ways to reduce this computation time moving forward. First, the signal could be downsampled prior to applying sliding-window spectral parameterization. In this paper, we wanted to match the instantaneous alpha total power estimation from the filter-Hilbert approach as closely as possible to ensure a fair comparison. So, we performed no downsampling. Because there is substantial overlap in the windows used to compute spectral features from time point to time point, such time resolution is unnecessary. The computation time would be reduced by the decimation factor chosen, greatly reducing computation time. Future studies could also investigate combining independent sliding-window estimations of alpha total power (through a filter-Hilbert approach) and of the aperiodic exponent.

In addition to the computational costs, there are fundamental questions surrounding how to identify oscillations. Although we took a spectral parameterization approach, there are several emerging methods for separating oscillations from aperiodic activity in both the time (Kosciessa et al., 2020; Seymour, Alexander, and Maguire, 2022) and frequency domains (Wen and Liu, 2016). Furthermore, there are more fundamental questions about how sensitive our methods are for even identifying oscillations in the first place (Van Bree et al., 2025), though there are methods for enhancing the oscillation signal-to-noise (Schaworonkow and Voytek, 2021; Schaworonkow and Nikulin, 2022). Finally, this analysis was constrained to looking at alpha power, though there are likely significant effects of alpha phase in working memory (Ai and Ro, 2014; Freunberger et al., 2009; Tran, Hoffner, et al., 2016), and potentially even with regards to the subtle differences in the alpha waveform (Bender, Voytek, and Schaworonkow, 2023; Cole and Voytek, 2017).

In terms of the kinds of tasks analyzed, this composite dataset worked well to investigate whether adjusting for aperiodic activity in estimating instantaneous alpha power substantially improved reconstruction of spatial location information in WM across an expansive number of WM tasks and participants. This investigation revealed an interesting contribution of aperiodic activity in representing spatial location during encoding, but the tasks were not designed to explicitly test for aperiodic activity and its role in encoding. Various modifications to the experimental task would allow for a more thorough examination of aperiodic activity’s role in WM encoding. First, using multiple stimuli and a pre-cue that identifies which stimulus is the target (i.e., should then be attended to) and which stimulus is the distractor (i.e., should not be attended to) would allow researchers to disambiguate whether aperiodic activity tracks attention-agnostic, stimulus-driven activity or attentive perception. Second, stimulus presentation was 250 ms or less in six of the seven tasks we analyzed; longer encoding periods would improve time-frequency analyses and ensure more of the time-frequency decomposition is computed with windows that are exclusively from the encoding period. Recent work by Tsubomi and colleagues demonstrated WM content is spontaneously removed after it becomes obsolete, suggesting WM is goal-directed and sensitive to task timing (Tsubomi et al., 2024). Manipulation of the length of the encoding period would allow for a similar test of sensitivity to task timing for the aperiodic exponent. Third, a WM task with serial presentation (e.g., Sternberg task) would allow researchers to evaluate whether each memory item is successively encoded by aperiodic activity since the corresponding neural activity will be separated in time. This would complement previous work that has suggested that sequential and simultaneous presentations rely on similar WM processes. For example, Woodman and colleagues showed that sequential and simultaneous presentation resulted in similar WM accuracy across different set sizes (Woodman, Vogel, and Luck, 2012), while Zhao and Vogel showed that performance is highly correlated in sequential and simultaneous WM tasks across individuals (Zhao and Vogel, 2023). Finally, similar to work done by Waschke and colleagues (Waschke, Wöstmann, and Obleser, 2017), researchers could experimentally manipulate the stimuli to bias aperiodic activity in the visual cortex and examine how such biasing affects WM encoding. Such an experiment would also enable explicit testing of the interaction between stimulus encoding by aperiodic activity and WM maintenance by alpha oscillatory activity.

## Conclusion

These results affirm the importance of alpha oscillations in maintaining spatial location information in WM, highlighting that it really is the oscillations’ power that is most relevant, not the total power. We developed novel methods for sliding-window estimation of alpha oscillatory power and aperiodic activity. These novel methods enabled us to disambiguate distinct roles for alpha oscillations and aperiodic activity in WM that had previously been mixed in the alpha total power signal. Specifically, aperiodic activity encodes spatial location during stimulus presentation, while the power of alpha oscillations maintain spatial location information throughout the WM delay. By leveraging sliding-window spectral parameterization, new experiments designed to test explicitly for the role of aperiodic activity in encoding could provide further mechanistic insights into how these distinct neural components contribute to WM encoding and maintenance. While the poor spatial resolution of scalp EEG may be masking differences in cortical processing of aperiodic and oscillatory activity, one implication of our results is that incoming visual sensory information drives cortical excitation in a topographic manner, but that this asynchronous, aperiodic drive is then translated into an oscillatory code for maintenance. Whether such an interaction between oscillations and periodic activity underlie encoding and maintenance of information in visual WM remains an open question.

## Data and code availability

All code used for all analyses and plots are publicly available on GitHub at https://github.com/voytekresearch.

## Author contributions

A.B., E.V., E.A., and B.V. conceived of the experiments. A.B. and B.V. developed the analyses. A.B. wrote the analysis code and analyzed the data. C.Z., E.V., and E.A. provided data and advised analyses. A.B. and B.V. wrote the manuscript, and all authors edited the manuscript.

## Acknowledgements

We thank Thomas Donoghue, Ryan Hammonds, Michael Preston, Natalie Schaworonkow, and John Serences for their thoughtful advice and feedback on the manuscript.

